# Structural and functional characterization of NEMO cleavage by SARS-CoV-2 3CLpro

**DOI:** 10.1101/2021.11.11.468228

**Authors:** Mikhail Ali Hameedi, Erica T. Prates, Michael R. Garvin, Irimpan Mathews, B Kirtley Amos, Omar Demerdash, Mark Bechthold, Mamta Iyer, Simin Rahighi, Daniel W. Kneller, Andrey Kovalevsky, Stephan Irle, Van-Quan Vuong, Julie C. Mitchell, Audrey Labbe, Stephanie Galanie, Soichi Wakatsuki, Daniel Jacobson

## Abstract

In addition to its essential role in viral polyprotein processing, the SARS-CoV-2 3C-like (3CLpro) protease can cleave human immune signaling proteins, like NF-κB Essential Modulator (NEMO) and deregulate the host immune response. Here, *in vitro* assays show that SARS-CoV-2 3CLpro cleaves NEMO with fine-tuned efficiency. Analysis of the 2.14 Å resolution crystal structure of 3CLpro C145S bound to NEMO_226-235_ reveals subsites that tolerate a range of viral and host substrates through main chain hydrogen bonds while also enforcing specificity using side chain hydrogen bonds and hydrophobic contacts. Machine learning- and physics-based computational methods predict that variation in key binding residues of 3CLpro- NEMO helps explain the high fitness of SARS-CoV-2 in humans. We posit that cleavage of NEMO is an important piece of information to be accounted for in the pathology of COVID-19.

## 1. Introduction

The impacts of the global Severe Acute Respiratory Syndrome Coronavirus-2 (SARS-CoV-2) pandemic have been extremely harsh. As of July 2021, SARS-CoV-2 has caused over 187 million confirmed cases of COVID-19, more than 4 million deaths (covid19.who.int/), and the global economy to contract by 3.5% in 2020 ^1^. Unlike previous *betacoronavirus* outbreaks, SARS-CoV-2 has spread to every country, which has provided the urgency and impetus to develop and rapidly distribute novel therapeutics to reduce the spread of the virus, including RNA-based vaccines. However, many societal impediments and the emergence of highly fit variants of concern are preventing vaccination coverage from reaching the levels necessary for herd immunity. Therefore, multiple therapeutic approaches will need to be implemented if we are to eventually eradicate SARS-CoV-2 and future zoonotic outbreaks.

One effective means to inhibit viral transmission and reduce the severity of the disease is to disrupt the lifecycle of the pathogen. SARS-CoV-2 is an enveloped *betacoronavirus* with a single stranded, positive-sense, 29 kb RNA genome that encodes a number of open reading frames (ORFs). *ORF1a* and *ORF1b* encode the polyproteins that are processed to generate the 15 nonstructural proteins (nsps) of SARS-CoV-2. These include the papain-like protease (PLpro) and the 3C-like protease (3CLpro), which are required to execute the viral life cycle and inhibit the host immune response^23^. These proteases therefore represent high-value targets for intervention.

The Protein Data Bank (RCSB PDB) is replete with structures of SARS-CoV-2 wild-type (WT) 3CLpro ^4, 5^. 3CLpro WT homodimers are a 67.60 kDa, heart-shaped complex ^4, 6–8^. Each 3CLpro chain consists of three domains. Domain I (aa. 8-101) and Domain II (aa. 102-184) have a predominantly β-sheet structure, form the active site, and contribute to dimerization. Domain III (aa. 201-303) is substantially α-helical and is the primary determinant of dimerization^6, 7, 9^. The active site of WT 3CLpro contains a catalytic dyad of His41 and Cys145, and an oxyanion hole formed by the main chain amide groups of Gly143 and Cys145^10^. During catalysis, His41 deprotonates the γ-thiol group of Cys145 to generate a nucleophile. Nucleophilic attack at the main chain carbonyl carbon of the P1 residue (immediately preceding the substrate scissile bond) forms a tetrahedral oxyanion intermediate. Heterolytic fission of the scissile bond stabilizes this intermediate, forms the acyl-enzyme intermediate, and releases the peptide downstream of the scissile bond. Finally, the active site is regenerated as a catalytic water molecule deacylates the acyl-enzyme intermediate.

Functionally, 3CLpro recognizes a hydrophobic substrate residue at P2 (usually Phe or Leu), a Gln at P1, and Ser, Val, Asn, or Ala residues at P1’. This recognition motif is found in multiple sites of the viral polyproteins, which are cleaved by 3CLpro to form mature nsp5-16. This consensus sequence is also present in proteins of the host innate immune pathway and therefore 3CLpro may blunt the immediate antiviral immune response via proteolysis ^11^. The NF-κB essential modulator (NEMO) ^12^ is one of the immune proteins that may be cleaved by 3CLpro. NEMO is necessary for activating NF-κB during the canonical NF-κB response signalling pathway, which is a critical first response to viral infection. A disrupted NF-κB pathway is a hallmark of chronic inflammatory diseases ^13^, which suggests that hNEMO cleavage by 3CLpro and the downstream dysregulation of NF-κB could contribute to the enhanced inflammatory response observed in COVID-19 patients ^12^.

The structure of 3CLpro from SARS-CoV-2 in complex with a NEMO-derived heptapeptide substrate was solved for the Porcine Epidemic Diarrhea Virus, an *alphacoronavirus* (PDB ID: 5ZQG)^14^. It is not known if this interaction is conserved in SARS-CoV-2 and other *betacoronaviruses.* In this work, *in vitro* assays showed that WT SARS-CoV-2 3CLpro can cleave the 33-residue peptide substrate, NEMO_215-247_. To explore the molecular basis of this interaction, we solved a 2.14 Å crystal structure of a SARS-CoV-2 3CLpro active site cysteine variant, C145S, in complex with the human decapeptide substrate NEMO_226-235_. This represents the first structure of SARS-CoV-2 3CLpro that is bound to a human peptide substrate. Using this structure as a starting point, extensive molecular dynamics simulations, quantum mechanics calculations, and machine learning-based predictions indicated that the few differences in NEMO and 3CLpro across host species and human-infecting *betacoronaviruses* significantly change the stability of the complex formed between these proteins. Finally, we discuss how ablation of NEMO via proteolysis connects with COVID-19 as a systemic disease.

## 2. Results

### 2.1. SARS-CoV-2 cleaves NEMO

In prior studies^11, 12^, SARS-CoV-2 3CLpro has been found to disrupt several components of the type I interferon pathway. NEMO has been shown to be cleaved by 3CLPro from feline infectious peritonitis virus (FIPV) and porcine epidemic diarrhea virus (PEDV) ^15, 16^. Molecular docking and comparative structural analyses suggest that NEMO similarly binds to SARS-CoV- 2 3CLPro in a position that favors proteolysis ^17^. Recently, N-terminomics experiments confirmed *in vitro* the cleavage of NEMO by SARS-CoV-2 3CLpro ^11^.

NEMO is known to form a homodimer of two, 419 residue (49 kDa), mostly ɑ-helical protomers ^18, 19^ and the overall domain architecture is well characterized (Figure 1a).^19^ A primary sequence analysis of NEMO (hNEMO) identifies five likely 3CLpro recognition sites around Gln83, Gln205, Gln231, Gln304, and Gln313, where the listed Gln would act as the P1 residue in each case ^20^ (Supplementary Figure S1). 3CLpro enzyme was incubated with five constructs of NEMO from mouse, *Mus musculus*, (mNEMO; i.e., amino acids 96-250, 221-250, 221-339, GST-96-250, GST-221-250, and GST-221-339), and the products detected by SDS-PAGE were consistent with cleavage following Gln231 (Supplementary Figure S2). To assess the ability and efficiency of the main protease to cleave hNEMO, we expressed and purified hNEMO truncated to amino acids 215-247, which mostly contains the Hlx2 domain, where the Gln231 recognition site is located. A previously reported assay was adapted.^21^ Concentration-dependent NEMO cleavage at the site reported for other coronaviruses (aa. 231-232) was detected via high performance liquid chromatography-mass spectrometry (HPLC-MS, Figure 1b).

**Figure 1.**
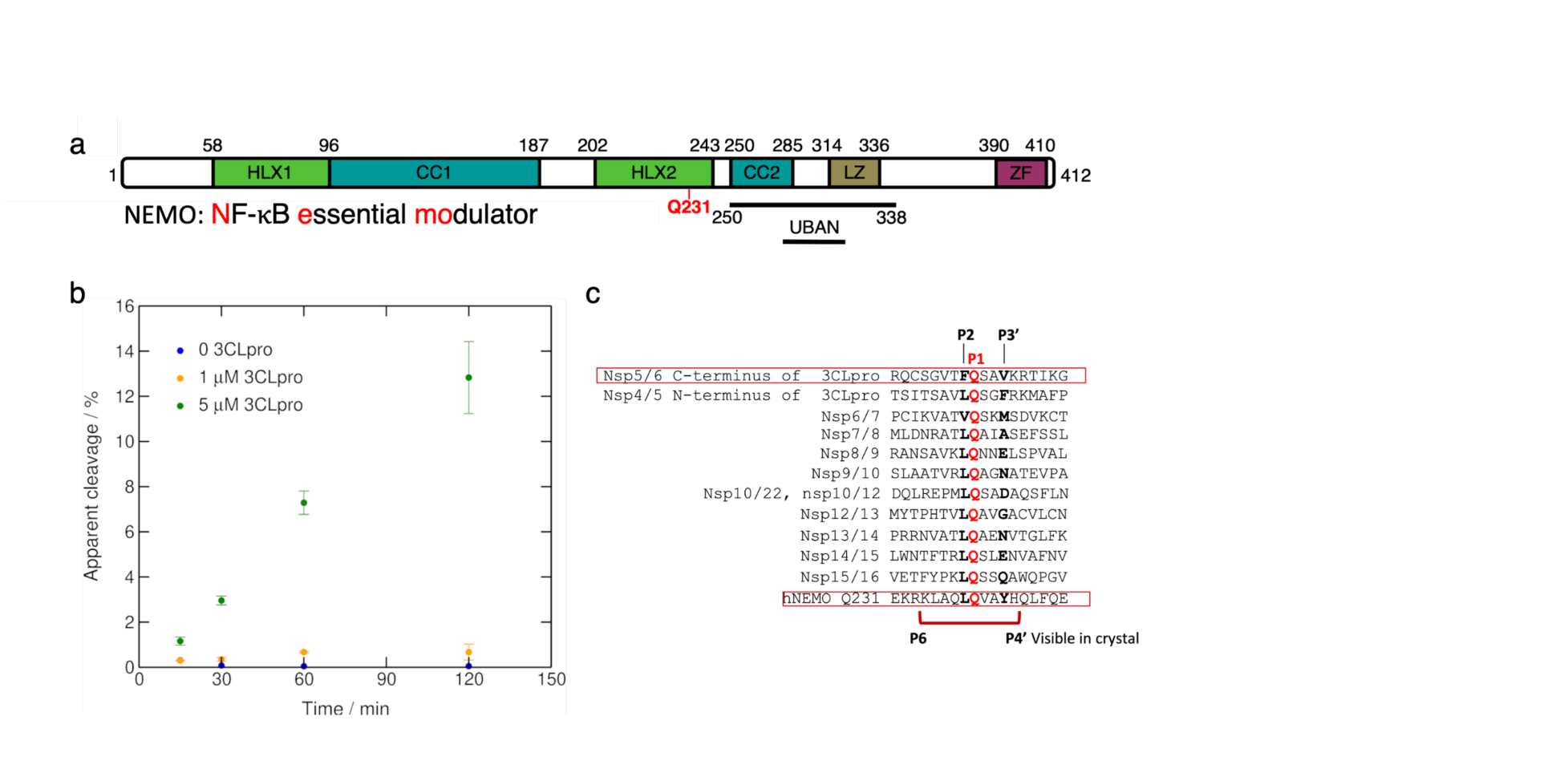
Truncated NF-κB essential modulator (NEMO) is cleaved by 3CLpro in enzymatic assays. **a)** NEMO includes the α-helical domain 1 (Hlx1), the coiled-coil domain 1 (CC1), the α-helical domain 2 (Hlx2), the coiled-coil domain (CC2), a leucine zipper (LZ) domain, and the C-terminal zinc-finger (ZF). Human NEMO (hNEMO) truncated at site 215 and 247 was used in enzymatic assays with 3CLpro. A recognition site of cleavage is found at Gln231. **b)** Cleavage of hNEMO_215-247_ at 0.053 μg/μL (∼13 μM) by 3CLpro at two concentrations. Reactions were incubated at 25 °C and aliquots were quenched at different times for analysis. The extent of proteolysis was quantified by LC-MS/MS. Apparent % cleavage was calculated by dividing the product peak area by the sum of the substrate and product peak areas. Error bars represent the range of duplicate enzymatic reactions. **c)** Multiple sequence alignment of peptide sequences of SARS-CoV-2 polyprotein and hNEMO. P1 site glutamine residues are shown in red. The region (P6 to P4’) visible in the crystal structure is indicated beneath the sequences.

### 2.2. Structure of 3CLpro C145S bound to a NEMO substrate peptide at 2.14 Å resolution

After extensive co-crystallization trials, we determined the structure of the SARS-CoV-2 3CLpro C145S variant in complex with a decapeptide from human NEMO at 2.14 Å resolution. This structure shows the Michaelis-like complex of SARS-CoV-2 3CLpro C145S and NEMO poised for catalysis.

In the NEMO-bound 3CLpro C145S structure, the asymmetric unit contains two, mature, heart- shaped 3CLpro C145S dimers. One dimer forms between chains A and B, and the second dimer forms between chains C and D. Chains B and C are each bound to the NEMO peptide (acetyl- KLAQLQVAYH-amide; aa. 226-235) (Figure 2a). Within the asymmetric unit, the C-terminal tail (Ser301-Gln306) of chain A swaps into the substrate-binding site of chain D. This connects the two dimers to each other in the asymmetric unit. Additionally, the substrate-binding site of chain A is bound by the C-terminal tail of chain D from a neighboring symmetry equivalent.

**Figure 2.**
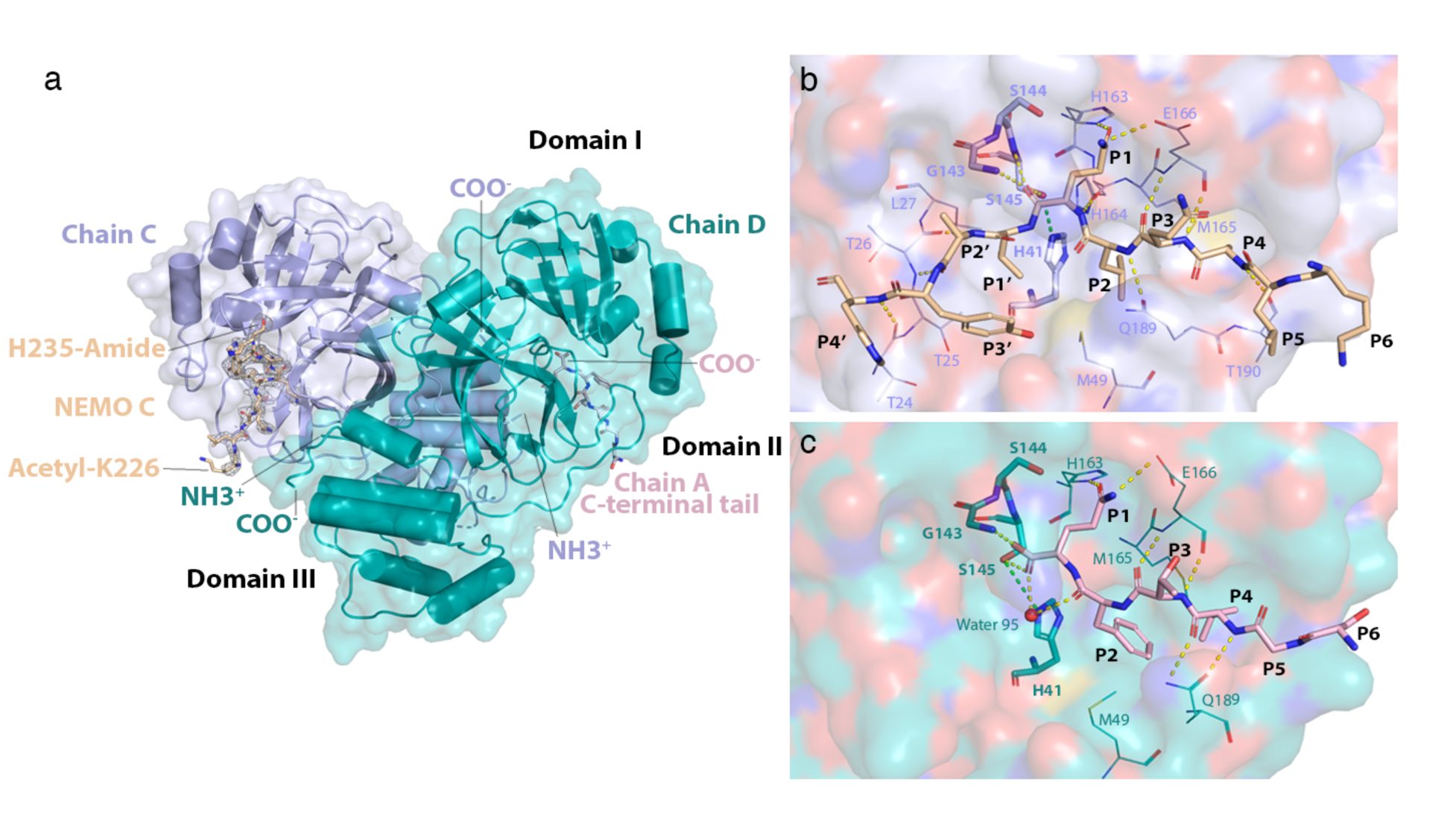
Structure of the NEMO-bound 3CLpro C145S homodimer. **a)** Global view of NEMO-bound 3CLpro dimer. Chains C and D have their N- (NH3+) and C-termini (COO-) labelled. NEMO bound into chain C (NEMO C) is displayed as sticks and colored wheat. A mesh of electron density is displayed around NEMO C in light grey. Acetylated Lys226 and amidated His235 at the N- and C-terminus of the NEMO peptide, respectively, are labelled. The C-terminal tail of chain A is colored in light pink, displayed as sticks and is observed to bind into the substrate-binding site of chain D. Domains I (aa. 8-101), II (aa. 102-184) and III (aa. 201-303) are labelled in black. **b)** Interactions of NEMO with the substrate-binding groove of 3CLpro C145S. NEMO is depicted in wheat. Residues Lys226 to His235 of NEMO are labelled P6 to P4’, respectively. The surface of chain C is shown in light blue and colored by atom type, where oxygens are red, nitrogens are dark blue, carbons are light blue, and sulfurs are yellow. Residues in the substrate-binding site of chain C are portrayed as lines and labelled. Catalytically relevant residues^22^ in chain C are portrayed as sticks and also labelled in bold. Hydrogen bonds are depicted as dashed yellow lines. The hydrogen bond that would form between Cys145 (in WT) or Ser145 (in C145S) and His41 is depicted as a dashed green line. **c)** Interactions of the C-terminal tail of chain A with the substrate-binding groove of 3CLpro C145S. The C-terminal tail of chain A is depicted as sticks in light pink. Residues Ser301 to Gln306 are labelled P6 to P1 respectively. The surface of chain D is shown in teal and colored by atom type. Residues in the substrate-binding groove of chain D that interact with the C-terminal tail of chain A are portrayed as lines and labelled. Catalytically relevant residues and hydrogen bonds are portrayed as in (b). The oxygen atom of water 95 is depicted as a red sphere.

This C-terminal tail-binding contributes to crystal lattice formation by connecting chains A and D to each other throughout the crystal (Supplementary Figure S3a). Finally, while the C-terminal tails of chains A and D are involved in forming the crystal lattice, the C-terminal tails of chains B and C bind at sites found at the interfaces of their respective dimers. Similar C-terminus swapping was reported for the C145A mutant of SARS-CoV-2 3CLpro ^10^. NEMO peptides bind in an extended conformation, similar to an anti-parallel β-sheet with 3CLpro residues Gln189-Ala191 and His163-Pro168. In chains B and C, there is density to model KLAQLQVAY (aa. 226-234) and, the entire peptide, KLAQLQVAYH (aa. 226-235), respectively (Supplementary Figure S3), where Lys226 is P6, His235 is P4’, Gln231 is P1 and Val232 is P1’ (Figure 2b). The N-acetyl and C-amide caps are not resolved. The densities in the substrate-binding sites of chains B and C can be unequivocally assigned as NEMO, rather than as a C-terminal tail. The densities do not connect to protein chains near the substrate-binding sites, but project into the bulk solvent. The densities in chains B and C are also too long to be assigned as the C-terminal tail, continuing beyond Ser145 in the substrate-binding site (unlike the density for the C-terminal tails in chains A and D). Finally, the positions of all C-terminal tails in the asymmetric unit are accounted for, and the shape of the density in chains B and C around residues P2, P3, P5, and P6 is consistent with the NEMO sequence.

### 2.3. Thr26 and Thr190 of 3CLpro pin the NEMO peptide into 3CLpro

Thr190 and Thr26 of 3CLpro use hydrogen bonds (H-bonds) to pin the ends of the NEMO peptide into the substrate-binding site, causing conformational changes in the site. Thr26 and Thr190 form H-bonds with Ala233 (P2’) and Ala228 of NEMO (P4), respectively. This causes the distance between the C_ɑ_ atoms of Thr26 and Thr190 to decrease from 21.7 Å (in WT 3CLpro) and 21.3 Å (in 3CLpro C145S without NEMO) to 20.4 Å in the NEMO-bound 3CLpro C145S.

### 2.4. Polar and hydrophobic subsites in 3CLpro position a glutamine at P1 while promoting substrate versatility

Our structure of 3CLpro C145S bound to a NEMO peptide identifies all the structural sites required for substrate-binding in 3CLpro. The substrate-binding site of 3CLpro consists of a groove made by four groups of consecutive residues, Gln189-Ala191, His163-Pro168, Phe140- Ser145, and Thr24-Leu27, as well as His41 and Met49. Gln189-Ala191 and His163-Pro168 run adjacent to each other, bind the P2-P4 residues of the NEMO peptide and position P1 in the active site. In the active site, Phe140-Ser145 bind P1. Following the active site, Thr24-Leu27 bind the P1’-P4’ residues. His41 is involved in catalysis and Met49 interacts with substrate P2. The subsites in the substrate-binding site of 3CLpro are identified as S4 to S4’, where S4 binds to P4 and S4’ binds to P4’ (Figure 2b).

Table 1 indicates that the preference for main chain H-bond interactions with the substrate retains substrate versatility in the S4-S2 and S1’-S4’ subsites. This supports the versatility of 3CLpro in cleaving the viral polyprotein at multiple sites and its activity towards several host proteins ^11, 12^. There are no S5 and S6 subsites in 3CLpro for P5 and P6 substrate residues, respectively.

**Table 1.** Interactions between SARS-CoV-2 3CLpro C145S and NEMO in the crystal structure.

| Subsite | 3CLpro residue (moiety) | Interaction (distance) | NEMO residue (moiety) |
| --- | --- | --- | --- |
| **S4** | Thr190 (main_C=O) | H-bond (2.7 Å) | Ala228 (main_N-H) |
|  | Met165 (side chain) | hydrophobic contact | Ala228 (side chain) |
| **S3** | Glu166 (main_C=O) | H-bond (3.0 Å) | Gln229 (main_N-H) |
|  | Glu166 (main_N-H) | H-bond (2.7 Å) | Gln229 (main_C=O) |
| **S2** | Gln189 (side_Oε1) | H-bond (2.9 Å) | Leu230 (main_N-H) |
|  | Met49 (side chain) | hydrophobic contact | Leu230 (side chain) |
|  | Met165 (side chain) | hydrophobic contact | Leu230 (side chain) |
| **S1** | His164 (main_C=O) | H-bond (2.8 Å) | Gln231 (main_N-H) |
|  | Ser145 (main_N-H) | H-bond (3.0 Å) | Gln231 (main_C=O) |
|  | Gly143 (main_N-H) | H-bond (3.1 Å) | Gln231 (main_C=O) |
|  | Ser145 ( γOH) | H-bond (2.8 Å) | Gln231 (main_C=O) |
|  | His163 (side_Nε2) | H-bond (2.4 Å) | Gln231 (side_Oε1) |
|  | Glu166 (side_Oε1) | H-bond (3.3 Å) | Gln231 (side_Nε2) |
| **S1'** | Thr25 (side chain) | hydrophobic contact | Val232 (side chain) |
|  | Leu27 (side chain) | hydrophobic contact | Val232 (side chain) |
| **S2'** | Thr26 (main_C=O) | H-bond (2.9 Å) | Ala233 (main_N-H) |
|  | Thr26 (main_N-H) | H-bond (2.9 Å) | Ala233 (main_C=O) |
| **S3'** | N.A. | N.A. | Tyr234 (N.A.) |
| **S4'** | Thr24 (main_C=O) | H-bond (2.6 Å) | His235 (main_N-H) |

To modulate substrate-selectivity, hydrophobic contacts select for hydrophobic side chains at P4, P2, and occasionally at P1’. Furthermore, in the S1 subsite, main chain interactions from His164 as well as Gly143 and Ser145 of the oxyanion hole combine with side chain interactions from His163 and Glu166 to select for glutamine at P1, position the main chain carbonyl carbon of NEMO Gln231 for the nucleophilic attack during catalysis, and stabilize the tetrahedral anion intermediate. Finally, the side chain of Ser145 forms H-bonds with the main chain carbonyl of Gln231, and Nε2 of His41 (3.0 Å). The H-bond between Cys145 and His41 is crucial in WT 3CLpro to generate the thiolate nucleophile for catalysis.

### 2.5. Comparison of bound NEMO and C-terminal tails helps clarify substrate-selectivity of 3CLpro

Comparison of NEMO and C-terminal-binding identifies that the S2 and S4 subsites contribute to substrate versatility in 3CLpro and that the S3 and S1 subsites are highly conserved between substrates. Compared to the C-terminal tail, NEMO forms an additional H-bond with Thr190 in the S4 subsite (Table 1). Additionally, NEMO forms an additional H-bond and hydrophobic contact with Gln189 and Met165, respectively, in the S2 subsite compared to the C-terminal tail. The S4 and S2 subsites therefore engage extra interactions to bind to NEMO P4 and P2 residues respectively. S3 and S1 subsites share the same interactions and so all the listed interactions are required for substrate-binding in 3CLpro. Finally, C-terminal P2 (Phe305) and P1 (Gln306) form H-bonds with the water 95 (3.0 Å and 2.7 Å respectively). This water is deacylating during 3CLpro catalysis ^10^, 3.3 Å from the C-terminal carboxyl carbon and has a Bürgi-Donitz angle of 122° (Figure 2c). In the NEMO-bound site, it is displaced by a hydrophobic side chain at P1’. Finally, the structure of 3CLpro bound to a C-terminal tail with residues in the S’ sites would identify such interactions in the S1’-S4’ subsites of 3CLpro.

### 2.6. C-terminal tails bind at two distinct sites of the 3CLpro dimer

An alternative binding site for the C-terminal tails is found in chains B and C in a groove formed at the dimer interface (Supplementary Figure S3a). At the dimer interface, the C-terminal tail is sandwiched between the two β-strands from Gly109-Tyr118 and Ser120-Arg131 in the neighboring chain and both the Ser1-Gly11 loop and Asp153-Cys156 turn in the same chain.

Specifically, in chain B, the side chain methyl group of Thr304 forms hydrophobic contacts with the side chain of Tyr118 in the neighboring chain A. Finally, the side chain of Phe305 forms hydrophobic contacts with a hydrophobic pocket consisting of the side chains of Phe8, Ile152, Val303, and the side chain propyl moiety of Arg298. It is possible that this interfacial site has a role in positioning substrates and cleavage products prior to and following catalysis, respectively.

### 2.7. Interfacial binding site of C-terminus attenuates self-inhibition of 3CLpro

In SARS-CoV 3CLpro, residues Arg298 and Gln299 in the C-terminal ɑ-helix are reported to be essential for dimerization and for activity by direct and indirect interactions with the S1 subsite of the neighboring 3CLpro in the dimer ^23^. In the C-terminal ɑ-helix of the NEMO-bound SARS-CoV-2 3CLpro C145S structure, Arg298 of chain B is closer to the carbonyl oxygen of Met6 when the C-terminal tail is bound at the dimer interface site than when it is bound in the substrate-binding groove. The closest distance of a side chain N in Arg298 from the backbone O in Met6 is 3.8, 3.6, 3.4, and 3.7 Å in chains A, B, C, and D, respectively, indicating tighter packing between the two sites with the C-terminal tail at the dimer interface. In both dimers within the asymmetric unit, this packing of Arg298 and Met6 is supported by a likely cation-π interaction between Arg298 and Phe8 as well as by H-bonds formed by Gln299 with the side chain of Ser139 in the neighboring protomer, chain A, and with the backbone of Arg4 in chain B. Molecular dynamics (MD) simulations of the 3CLpro dimer, using chains A and B, with NEMO bound to the latter, show that these interactions involving Arg298 and Gln299 persist during most of the trajectories except by the pair Arg298-Met6, indicating that other inter-dimeric contacts contribute to stabilizing the non-interfacial conformation of the C-terminus (Supplementary Figure S4). Finally, estimation of the binding affinity by rigid re-docking of a C-terminus-like peptide (Cys300-Gln306) to the two alternative C-terminal binding sites predicts stronger binding to the catalytic site (-14.5 kcal/mol) compared to the dimer interface (-12.8 kcal/mol) that by far compensates for a weaker Arg298-Met6 interaction. These comparable binding affinities suggest that the interfacial binding site contributes to decreasing self-inhibition of 3CLpro. In line with that and with the NEMO-cleavage detected *in vitro*, the interaction with C-terminal tail at the active groove is less effective compared to NEMO-binding (-19.8 kcal/mol).

### 2.8. Molecular dynamics simulations of the complex formed by WT 3CLpro and a long construct of NEMO

MD simulations performed on a longer construct of hNEMO with 3CLpro carrying catalytic Cys145 recapitulated the conformation captured experimentally (Figure 3a). A model of a partially unfolded hNEMO dimer (aa. 190-270) bound to 3CLpro was used in the simulations, which show a conformational fluctuation of the binding core of hNEMO (aa. 227-234) near the crystallographic conformation, with a C_ɑ_ RMSD of 3.2 ± 0.8 Å (Figure 3a). The distance between the catalytic S^-^ in Cys145 and the carbonyl C in Gln231 stands near 5.0 Å (5.0 ± 0.4 Å), indicating a conformation that would favor the nucleophilic attack by 3CLpro (Supplementary Figure S5).

**Figure 3.**
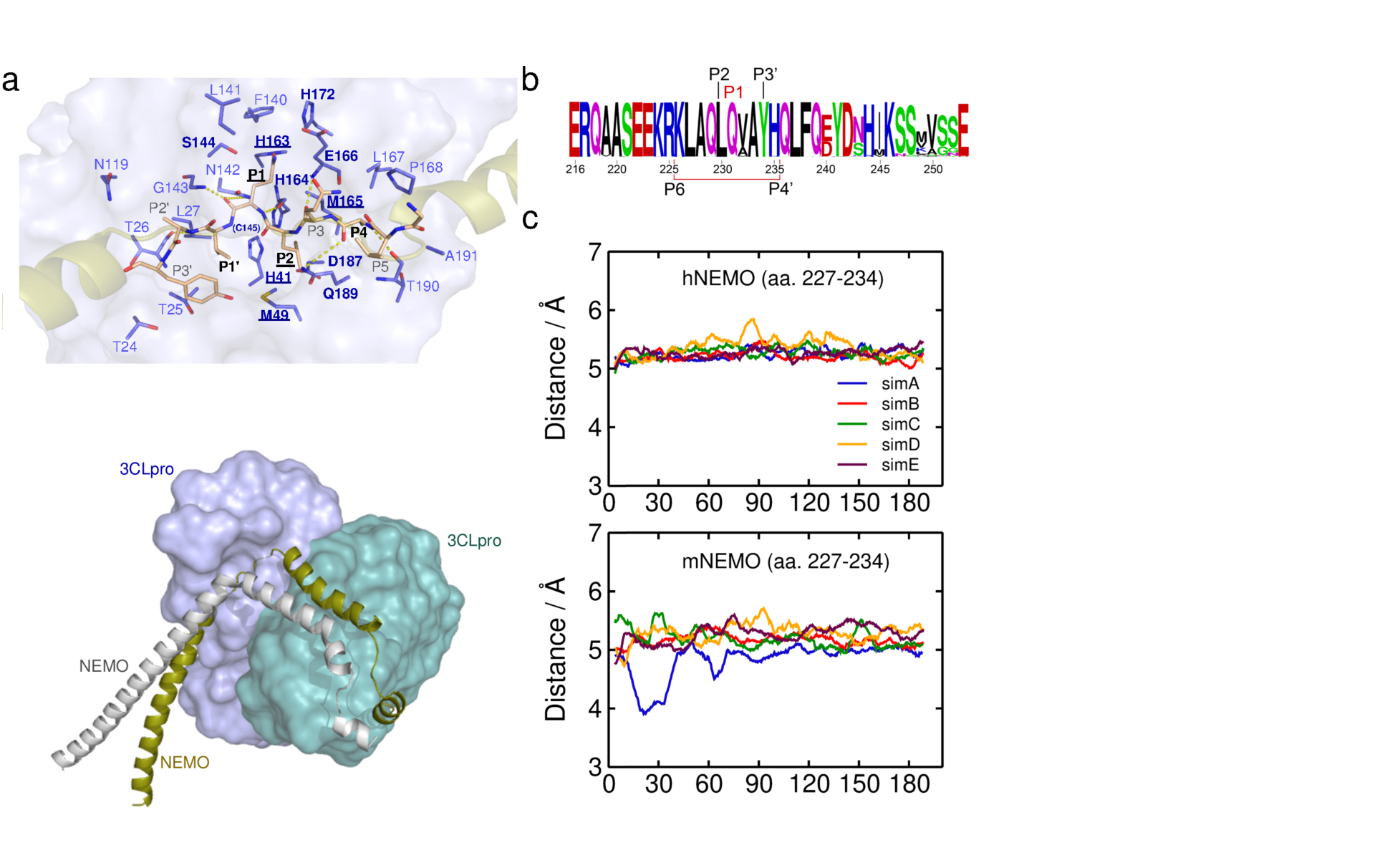
Molecular dynamics simulations of SARS-CoV-2 3CLpro bound to human or mouse NEMO. a) Main contacts from MD simulations and predicted hot spots in WT 3CLpro bound to human NEMO. Contacts that persist for more than 70% of the simulation time are depicted. Hot spots predicted with either KFCa or KFCb are labeled in bold and those predicted with both methods are underlined. Persistent H-bonds are shown as dashed lines (Supplementary Table S1). A representative conformation of the short (aa. 227-234) and long (aa. 190-270) constructs of NEMO in the binding site is shown as solid and transparent surfaces, respectively. The lower panel displays a MD snapshot of the longer NEMO dimer partially unfolded, with one protomer bound to 3CLpro. **b)** Sequence Logo generated with WebLogo ^24^ of NEMO (aa. 216-253) across 535 animal sequences, including (but not limited to): placentals, bats, marsupials, birds, rodents, and primates. Clustal Omega ^25^ was used for multiple sequence alignment. **c)** Time evolution of the distance between the catalytic S^-^ in 3CLpro Cys145 and the carbonyl C of Gln231 in human NEMO (hNEMO) and mouse NEMO (mNEMO) computed from MD simulations.

Simulations of the WT 3CLpro dimer (chains A and B) bound to hNEMO_227-234_, using our crystal structure as the initial configuration, show that the H-bonds identified in the static structure persist (Supplementary Table S1, Figure 3a). Additional H-bonds are formed between the residue pairs Ala228-Gln189, Gln231-Cys145, and Val232-Asn142, in hNEMO and 3CLpro, respectively, appearing during about 20-30% of the simulation time. The H-bond pair Leu230- Gln189 appears less than 7% of the time. Several other residues in the binding site of 3CLpro form stable contacts with hNEMO (Figure 3a). Finally, using the crystal structure as input (chain B, NEMO B), the machine learning (ML)-based predictor, KFC2^26, 27^ indicates that hot spots, i.e., residues likely accounting for most of the binding affinity, coincide with those forming persistent contacts between 3CLpro and hNEMO (Figure 3a).

### 2.9. Predicted binding affinity of NEMO to 3CLpro differ among host species and coronaviruses

The binding core of NEMO is highly conserved among different species (Figure 3b), but amongst mammals, the predicted hot spot at P1’ is a Val232 in hNEMO whereas it is an Ala232 in mNEMO and golden hamster (*Mesocricetus auratus*) NEMO. Next, we carried out other MD simulations for hNEMO_227-234_ and mNEMO_227-234_. Like in hNEMO, the distance between the catalytic S^-^ in 3CLpro Cys145 and the carbonyl C in mNEMO Gln231 remains near 5.0 Å (Figure 3c). However, the simulations show a decrease in the average number of contacts between mNEMO (49 ± 4) and 3CLpro compared to hNEMO (54 ± 4), and the C_ɑ_ root-mean-square deviation (RMSD) of mNEMO (1.9 ± 0.6 Å) relative to the initial position is larger than hNEMO (1.4 ± 0.4 Å). The impact of V232A becomes more pronounced in simulations with NEMO_190-270_ in the dimeric form, in which we observe large fluctuations in the distance between 3CLpro Cys145 and mNEMO Gln231 (Supplementary Figure S5). This is explained by competing interactions within the NEMO dimer. NEMO dimerizes at Hlx2 forming hydrophobic contacts between the pairs of Tyr234, Leu230, and Leu227 in each protomer ^18^, and Tyr234 forms H-bonds with Glu240 in the neighboring protomer.

The viral counterpart, 3CLpro, is also highly conserved among species, but structural differences among human-infecting *betacoronaviruses* underpin differences in their predicted interactions with NEMO. Specifically, in HCoV-HKU1, MERS-CoV, SARS-CoV, and SARS-CoV-2, the predicted hot spots, Met49, His164, and Met165, are not fully conserved, and the loop/ɑ-helix formed by aa. 41-53 in Domain I of 3CLpro, which skirts the substrate-binding site, harbors most of the other differences. MD simulations of these proteins bound to hNEMO (aa. 227-234) show that 3CLpro from MERS-CoV (43 ± 4) and SARS-CoV (46 ± 7) exhibit lower average number of contacts than SARS-CoV-2 (52 ± 3) and HCoV-HKU1 (53 ± 2) (Figure 4a). Particularly, this corroborates the hypothesis, detailed in the Discussion, that the S46A substitution between SARS-CoV-2 and SARS-CoV 3CLpro significantly impacts interactions with NEMO. Unsurprisingly, given the proximity of S/A46, the hot spot Met49 is one of the affected contacts. Simulations of these enzymes in the *apo* state show that the aa. 41-53 in 3CLpro exhibit slightly different flexibility, with SARS-CoV 3CLpro displaying a RMS fluctuation of 2.2 and 2.3 Å in each chain of the dimer and SARS-CoV-2 3CLpro, 2.3 and 2.5 Å. Additionally, essential C_ɑ_ cross-correlation analysis suggests that SARS-CoV-2 3CLpro dimer has significantly more residue pairs exhibiting synchronized motions along the same or opposite directions than SARS-CoV 3CLpro, which may reflect a tighter dimeric packing (38183 *vs.* 22513 pairs and 35707 *vs.* 17567 pairs, respectively; cutoff is a correlation modulus of 0.85; Supplementary Figure S6).

**Figure 4.**
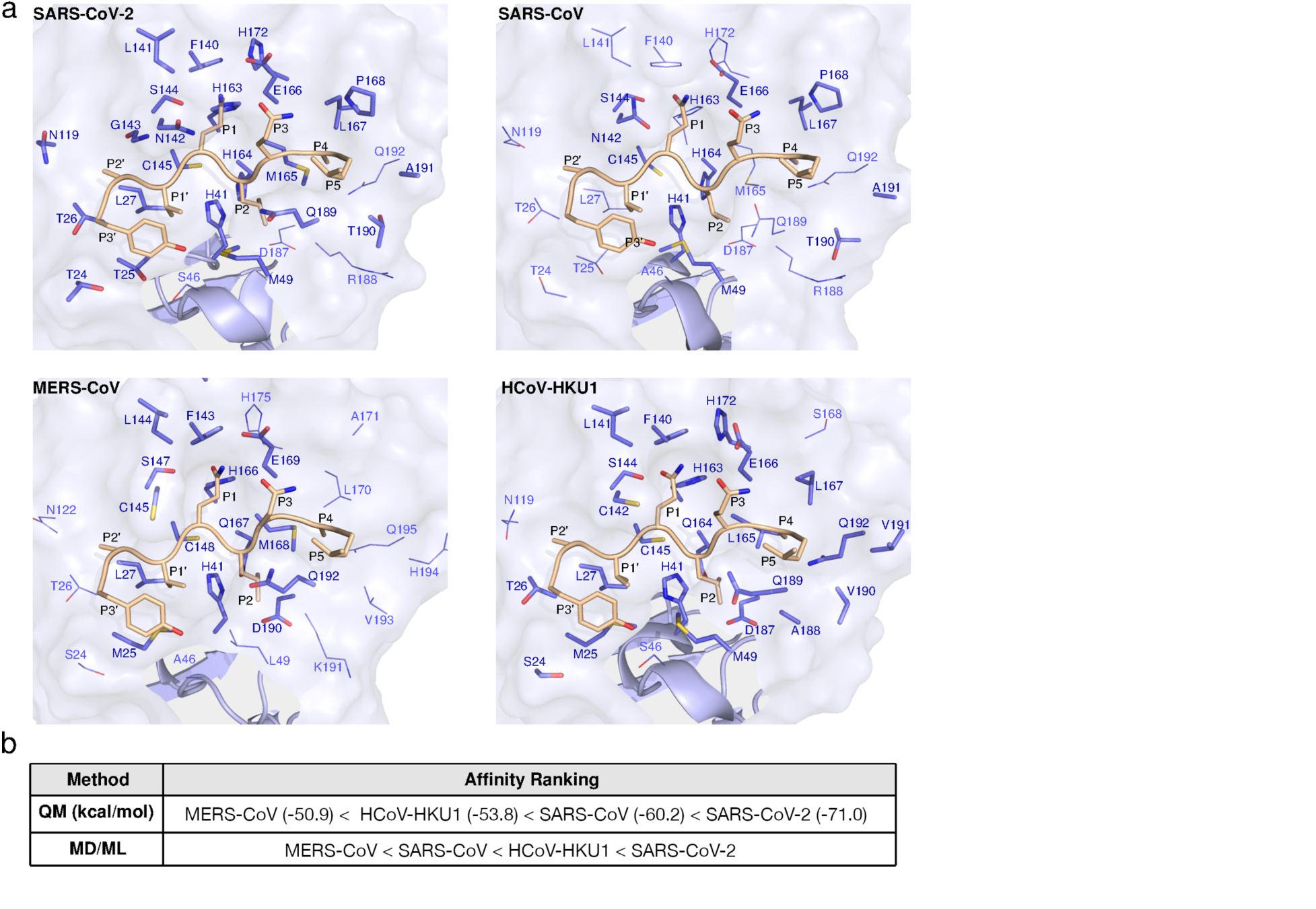
Comparative analysis of 3CLpro from human-infecting *betacoronaviruses* bound to NEMO. a) Substrate-binding site of human-infecting *betacoronaviruses* with hNEMO_227-234_. Contacts that persist for more than 70% of the simulation time are labeled in bold. **b)** Ranking of predicted NEMO-binding affinities to 3CLpro from *betacoronaviruses* computed with quantum mechanics (QM, values of internal binding energy as defined in the Method section are shown in parentheses) and molecular dynamics/machine learning (MD/ML) approaches. In the MD/ML approach, five machine learning methods were used to train the model, namely, support-vector machine, gradient-boosted trees (scaled and unscaled*), and random forest (scaled and unscaled*). The ranking displayed was consistent for nine of the ten cases of MD conformers (5 ML models applied to either all MD conformers or the three lowest-energy MD conformers, yielding 5 x 2 = 10 cases). The exception was a boosted tree model trained on unnormalized features that yielded a ranking of SARS-CoV < MERS-CoV < HKU1-CoV < SARS-CoV-2 when considering just the conformers with the lowest energy of interaction with 3CLpro computed from MD simulations. *Unscaled refers to the fact that unscaled, or unnormalized, features were used in the training.

Finally, for a quantitative assessment of relative hNEMO-binding affinities, quantum mechanics (QM)- and MD/ML-based predictions were carried out. In the QM calculations, the fragment molecular orbital density-functional tight-binding (FMO-DFTB) method was used. For the MD/ML-based approach, five different models were trained. The rankings from the weakest to the strongest binding enzyme were nearly consistent between the two approaches (Figure 4b).

The exception is the relative position of SARS-CoV 3CLpro. A possible explanation, further explored in Discussion, is that the few differences between SARS-CoV and SARS-CoV-2 3CLpro affect the conformational flexibility of the binding site, which may be captured in MD/ML but not in the QM approach.

## 3. Discussion

### Predicted mechanism of NEMO-binding to SARS-CoV-2 3CLpro

SARS-CoV-2 3CLpro binds Lys226-His235 of the α-helical Hlx2 domain of one NEMO protomer in an extended conformation. In agreement, QM calculations predict favorable energetics for NEMO-binding to 3CLpro in such an extended conformation. SARS-CoV-2 3CLpro therefore either makes use of a transient partially unwound state of Hlx2 for proteolysis or it actively outcompetes NEMO dimer interactions and unwinds the α-helix of one protomer. Specifically, 3CLpro forms two hydrophobic contacts as well as H-bonds with Leu230. This outcompetes the hydrophobic contacts formed by Leu230 in the NEMO dimer partner.

Additionally, 3CLpro forms H-bonds with NEMO His235 and Ala233, either side of Tyr234, outcompeting the H-bond formed between Tyr234 and Glu240 in the NEMO dimer.

### Functional impact of NEMO proteolysis

NEMO homodimers form three cellular structures. First, NEMO homodimers. Second, a higher-order lattice ^28^. The lattice assembles through both non-covalent associations of the IKKα/β domains of NEMO homodimers and polyubiquitin-binding between NEMO homodimers. Third, NEMO lattices compact to form signalosome-proximal aggregates during NF-κB signaling stimulation by interleukin-1 (IL-1) ^28^. Higher-order structures cooperatively enhance NF-κB signaling. Cleavage at Gln231 in the NEMO homodimer would separate the NEMO kinase-binding site from the ubiquitylation and ubiquitin-binding sites, preventing IKKα/β activation, IKK assembly, and thus lattice formation, which all ultimately ablate NF-κB signaling. It is unknown if and to what extent Gln231 is accessible to 3CLpro in the NEMO aggregates. Future characterization of the different structural levels of NEMO as well as how they interchange in equilibrium can help to answer this question.

### Dissecting the versatile substrate-selectivity of 3CLpro using peptide-bound structures

Comparison of the NEMO-bound structure with the structure of SARS-CoV-2 3CLpro C145A bound to a peptide of its N-terminal autoprocessing sequence, acetyl-SAVLQSGF-amide (PDB ID: 7N89) ^30^ identifies specific interactions engaged by 3CLpro to bind either the N-terminal or NEMO substrate (Figure 5), as well as conserved interactions between the two structures that are required for substrate-binding.

**Figure 5.**
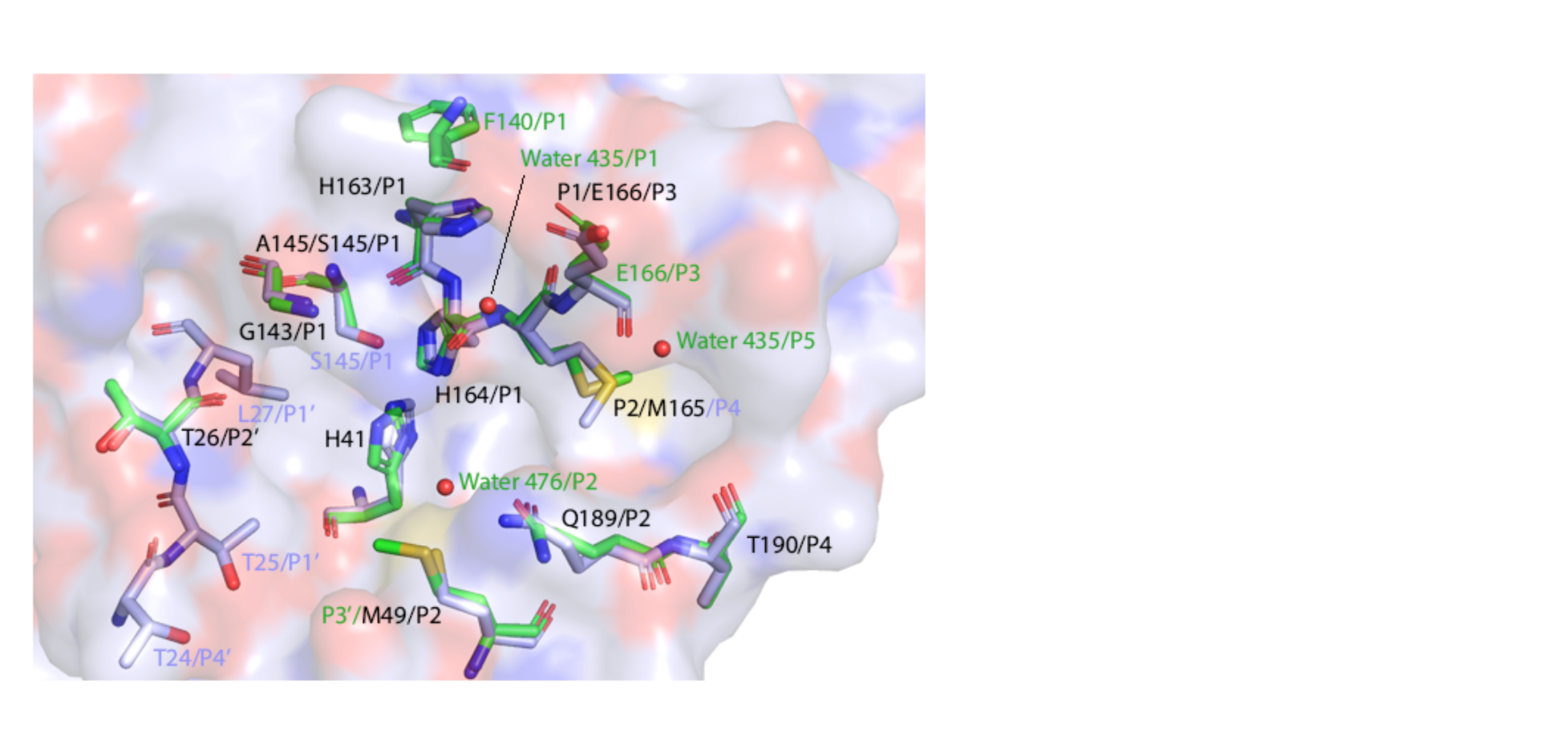
Comparison of 3CLpro interactions with its N-terminal sequence and NEMO. The surface of chain C from our NEMO-bound structure is depicted as in Figure 2b. This is juxtaposed with residues from our structure that form interactions with NEMO (light blue sticks) as well as residues (green sticks) from a structure of 3CLpro bound to an N-terminal peptide (PDB ID: 7N89). Residues forming conserved interactions are labelled in black. Those found only in NEMO-bound 3CLpro are labelled in light blue. Those found only in 7N89 are labelled in green. Water molecules are portrayed as red spheres and are exclusive to 7N89. Met165 forms a hydrophobic contact with the P2 and P4 residues of NEMO but only with P2 with the N-terminal peptide. Met49 forms a hydrophobic contact with the P2 residues of both NEMO and the N-terminus, but also forms hydrophobic contacts with P1’ and P3’ of the N-terminal sequence. Substrates are excluded for clarity.

In 7N89, the S5 subsite recruits a water molecule to bind N-terminal P5 (Ser). This is not observed in our NEMO-bound structure. The S4 subsite forms a hydrophobic contact between Met165 and NEMO P4 but not with N-terminal P4 (Ala). This site therefore contributes to the substrate-versatility of 3CLpro. In the S3 subsite, the main chain H-bonds from Glu166 are conserved when binding both substrates. However, 3CLpro recruits an additional hydrophobic contact between the alkyl moiety of Glu166 and N-terminal P3 (Val). The S2 subsite conserves all interactions observed between 3CLpro and NEMO but engages an additional main chain H-bond with water476 that stabilizes N-terminal P2 (Leu) in its subsite. 7N89 identifies extra interactions that strengthen the specificity for Gln P1. In 7N89, the side chain of P1 Gln forms additional H-bonds with the main chain carbonyl of Phe140 and water472. Due to the polar nature of the Ser145 γ-hydroxyl group in 3CLpro C145S, there is an additional observed H-bond with the carbonyl group of Gln P1 in the NEMO-bound structure. The S1’ subsite is flexible. It forms no interactions with the N-terminal substrate but engages hydrophobic contacts from Thr25 and Leu27 side chains to bind NEMO. It therefore contributes to 3CLpro substrate versatility. The S2’ subsite interactions are conserved between NEMO- and N-terminal-bound structures, indicating a requirement for Thr26 peptide-nonspecific main chain interactions at this subsite. Finally, the S3’ and S4’ subsites both contribute to substrate versatility. 3CLpro engages hydrophobic contacts between Met49 and Thr24 to bind N-terminal P3’ (Phe) and NEMO P4’ respectively.

### Differences in betacoronaviruses 3CLpro and in host NEMO likely affect pathogenesis

Subtle structural differences between otherwise highly conserved *betacoronaviruses* 3CLpro enzymes are predicted to have substantial effects on NEMO-binding. Our computational analyses suggest that the single substitution S46A in the substrate-binding site of SARS-CoV-2 3CLpro relative to SARS-CoV-2 3CLpro increases local rigidity and, with that, conformational changes that optimize the interactions with NEMO are impaired^31^. Alanine has significantly higher ɑ-helical propensity than serine^32^ and, indeed, Ala46 is part of an ɑ-helix while Ser46 is at a turn in *apo* SARS-CoV^33^ and SARS-CoV-2 3CLpro, respectively. MERS-CoV 3CLpro has Ala46 and Pro45, both known to increase local rigidity, ^34^ and HCoV-HKU1 lacks such rigidifying residues. This is consistent with the predicted decrease in NEMO affinity from SARS-CoV-2 through HCoV-HKU1, SARS-CoV, and MERS-CoV. This may also explain why N-terminomics identified more substrates for SARS-CoV-2 3CLpro than for SARS-CoV 3CLpro despite 96% identity ^11^ and why FRET experiments did not capture any activity of SARS-CoV 3CLpro on NEMO^16^.

Differences distal to the binding site may also have a significant functional impact. For example, the presence of Thr285 in the Domain III of SARS-CoV 3CLpro is associated with a looser dimer and slightly lower catalytic efficiency compared to SARS-CoV-2 3CLpro, which has Ala285^8^. Essential dynamics analysis shows that this difference in dimerization was captured in our trajectories. Although a causal relationship is not obvious, our results reinforce the importance of accounting for molecular differences in the entire proteome for a complete understanding of pathogenesis.

Similarly, we predict that the substitution V232A in NEMO from human to mouse affects NEMO-binding to 3CLpro. Site 232 is variable in other relevant reservoirs of SARS-CoV-2 and may have implications in the disease tolerance (Supplementary Table S2)^35^. Therefore, animal models incorporating multiple elements of the human immune system, as in mice transplanted with human immune cells ^36^, may be a way of accounting for the impact of apparently subtle differences in pathogenesis.

### Highly fit SARS-CoV-2 can fine-tune viral replication and modulation of host immune responses

The interaction of 3CLpro with host proteins that are part of the innate immune response appears to be finely tuned to avoid competition with its role of processing the viral polyproteins and to maintain functional cells for production of virions ^37^. Our enzymatic assays indicate that cleavage of NEMO is a slow process and likely not impactful in the first hours of infection ^38^.

Similarly, it was demonstrated that SARS-CoV-2 infection of different cell lines requires 24 to 48 hours to cause a dramatic reduction in the levels of host proteins cleaved by PLpro and 3CLpro ^12^. These observations and the cleavage of multiple host proteins detected with N-terminomics suggest that the cleavage of NEMO is one component of a number of mechanisms that SARS-CoV-2 combines to counteract the host immune response ^11^. However, certain traits of COVID-19 are remarkably consistent with known effects caused by NEMO ablation, suggesting its particular relevance in the pathophysiology. Mutations in the *NEMO* gene in the genetic disease *incontinentia pigmenti* are typically lethal in males and cause skewed X-inactivation patterns in females, selecting for the normal allele, as *NEMO* resides on the X- chromosome ^39, 40^. Similarly, males seem to develop more severe COVID-19 ^41^. Additionally, deletion of *NEMO* leads to rarefaction of brain microvessels ^42^ and increased vascular permeability in the brain is observed in COVID-19 patients with neurological symptoms ^43^, which resembles those of *incontinentia pigmenti* ^44^.

### Concluding remarks

NEMO lies at the nexus of the antiviral response driven by the mitochondrial antiviral-signaling protein (MAVS) that results in the activation of both NF-κB and type I interferons^45^. The NF-κB pathway is targeted by diverse viruses ^46^ and its suppression contributes to an imbalance in the renin-angiotensin system, which is proposed to result in severe COVID-19 outcomes^47^. Our results suggest that SARS-CoV-2 weakens innate immunity by cleaving NEMO using 3CLpro.

In addition to directly countering the immunosuppressive effects from NEMO cleavage, inactivating 3CLpro as a therapeutic strategy would impair viral replication and reduce the production of proteins that intervene downstream from MAVS, such as nsp6, and nsp13 ^48, 49^.

Future avenues for research will involve defining the link between the binding affinity of 3CLpro to NEMO and the observed proteolytic rate via enzymatic assays. Finally, semi-quantitative analysis of accumulation of 3CLpro and the products of NEMO-cleavage in human cells from different tissues infected with SARS-CoV-2 will help to unravel the specific pathogenesis traits derived from ablation of NEMO.

## 4. Acknowledgements

We would like to acknowledge funding from DOE Office of Science through the National Virtual Biotechnology Laboratory, a consortium of DOE national laboratories focused on response to COVID-19, with funding provided by the Coronavirus CARES Act. This work was partially funded from the Laboratory Directed Research and Development Program of Oak Ridge National Laboratory, managed by UT-Battelle, LCC for the US Department of Energy, LOIS:10074, which supported the conceptual work on the NEMO cleavage in animal models for COVID-19 pathology. Funding for human pathogenesis conceptualization was provided by the National Institutes of Health grant 3RF1AG053303-01S2. Use of the Stanford Synchrotron Radiation Lightsource, SLAC National Accelerator Laboratory, was supported by the U.S. Department of Energy, Office of Science, Office of Basic Energy Sciences under Contract No. DE-AC02-76SF00515. The SSRL Structural Molecular Biology Program is supported by the DOE Office of Biological and Environmental Research, and by the National Institutes of Health, National Institute of General Medical Sciences (P30GM133894). The contents of this publication are solely the responsibility of the authors and do not necessarily represent the official views of NIGMS or NIH. We also thank Daniel Fernandez and the Macromolecular Structure Knowledge Center at Stanford for providing equipment for crystallography. This research used resources of the Oak Ridge Leadership Computing Facility, which is a DOE Office of Science User Facility supported under Contract DE-AC05-00OR22725. This research also used resources at the Spallation Neutron Source and the High Flux Isotope Reactor, which are DOE Office of Science User Facilities operated by the Oak Ridge National Laboratory. The Office of Biological and Environmental Research supported research at ORNL’s Center for Structural Molecular Biology, a DOE Office of Science User Facility.

## 5. Methods

### 5.1. Enzymatic assays

#### NEMO peptide expression and purification

The constructs of Human NEMO (residues 215-247) and mouse NEMO (residues 221-250) cloned into pGEX-6p-1 vector were transformed into BL21 (DE3) cells and selected using ampicillin-enriched LB media. Cells were grown to OD600 of 0.6-0.8 and induced with 0.25 mM IPTG overnight at 25°C. Cells were harvested, resuspended in phosphate buffered saline (PBS) and lysed on ice using sonicator (QSonica). Insoluble material was pelleted using centrifugation and the lysate was incubated with Glutathione Sepharose 4B (GS4B, Cytiva) for 2 hours on a rotating platform at 4°C. The beads with the GST-tagged protein were separated using a gravity column and washed with PBS to remove any unbound protein. The GST tag was cleaved on-column using Prescission protease and incubating overnight at 4 °C. Cleavage was confirmed by SDS-PAGE and the cleaved protein was eluted using PBS. The protein was further purified by size-exclusion chromatography on a Superdex 75 16/60 column using 20 mM Tris-HCl, pH 8.0 and 150 mM NaCl as the running buffer.

#### 3CLpro expression and purification

3CLpro WT enzyme for assays was prepared independently from a clone of the SARS-CoV-2 NSP5 gene in pD451-SR (Atum, Newark, CA) according to published procedure ^6^. To make the authentic N-terminus, the protease sequence is preceded by maltose binding protein followed by the 3CLpro autoprocessing site between NSP4 and NSP5 (SAVLQ↓SGFRK, arrow indicates the cleavage site). Authentic C-terminus is achieved by a sequence of a human rhinovirus 3C (HRV-3C) cleavage site (SGVTFQ↓GP) connected to a His6 tag. The N-terminal flanking sequence is autoprocessed during expression in *E. coli* (BL21 DE3), whereas the C-terminal flanking sequence is removed by the treatment with HRV-3C protease (Millipore Sigma, St. Louis, MO) in-between two rounds of Ni-NTA affinity chromatography.

#### Human NEMO cleavage assay

70 μL reactions of 0.053 μg/μL substrate hNEMO 215-247 and 0, 1, or 5 μM WT 3CLpro were prepared in duplicate using a reaction buffer of 20 mM Tris-HCl, pH 7.35, 100 mM sodium chloride, 1 mM EDTA, and 2 mM reduced glutathione. Reactions were incubated at 25 °C with gentle shaking and 5 μL aliquots were quenched by diluting into 95 μL of 1.63% formic acid in water at 4 °C at 15, 30, 60, and 120 mins. 2 μL of quenched reactions were injected onto an Agilent EclipsePlusC18 1.8 μM, 2.1 x 50 mm chromatography column, and eluted using a gradient elution of 2-80% buffer B (0.1% formic acid in acetonitrile) against buffer A (0.1% formic acid in water) over 8.5 mins. Samples were introduced into mass spectrometers, and molecular masses of the substrate [M+2H]+2 ion with m/z 1486.5 and the C-terminal fragment VAYHQLFQEYDNHIKS [M+H]+ ion with m/z 1675.5 were detected using positive mode ionization, a capillary voltage of 4000 V, a nozzle voltage of 1500 V, an MS2 scan of 830-1490 m/z over a 330 ms scan time, a fragmentor voltage of 300 V, and a cell accelerator voltage of 3 V. Substrate and product peak areas were determined by integrating the extracted ion chromatograms.

#### Mouse NEMO cleavage assay

NEMO peptides were prepared as 0.2 mg/mL stocks and 3CLpro as 0.5 *μ*M stock in the same assay buffer as described for the human NEMO cleavage assay, and assays were initiated by adding 5 *μ*L enzyme (or buffer for negative controls) to 5 *μ*L peptides. Reactions were incubated at 37 °C in a thermocycler for 30 min and quenched by adding 10 *μ*L quench buffer (50% v/v NuPAGE™ 4x LDS buffer, 20% v/v 0.5 M dithiothreitol, 30% v/v water) and heating at 37 °C in a thermocycler for 20 min. A NuPAGE™ 4 to 12%, Bis-Tris, 1.0 mm, Mini Protein Gel, 12- well, was loaded with 10 *μ*L/lane and 8 *μ*L SeeBluePlus2 ladder, and proteins were separated with 200 V electrophoresis for 35 min with MES buffer. Bands were visualized with BullDog Bio Acquastain.

### 5.2. Crystallography

#### 3CLpro WT expression and purification

BL21(DE3) cells were transformed with pMCSG53 pDNA containing a 3CLpro WT insert with an autoprocessing-sensitive N-terminal Maltose Binding Protein (MBP) tag and a PreScission protease-sensitive C-terminal His6 tag (provided by Andrzej Joachimiak). Transformants were selected using ampicillin-enriched LB media, grown to OD600 of 0.6-0.8 and induced over 10 hours with 0.5 mM IPTG (GoldBio, USA), and harvested. Cell pellets were resuspended in 50 mM HEPES pH 7.2, 150 mM NaCl, 5% glycerol, 20 mM Imidazole, 10 mM 2-mercaptoethanol, and lysed by sonication. Insoluble material was pelleted by centrifugation and the lysate loaded onto a Ni^2+^ column, equilibrated with 50 mM HEPES pH 7.2, 150 mM NaCl, 5% glycerol, 10 mM Imidazole, 10 mM 2-mercaptoethanol. The column was washed using 50 mM HEPES pH 7.2, 150 mM NaCl, 5% glycerol, 50 mM Imidazole, 10 mM 2-mercaptoethanol, and eluted using 50 mM HEPES pH 7.2, 150 mM NaCl, 5% glycerol, 500 mM Imidazole, 10 mM 2- mercaptoethanol. The C-terminal His6 tag was cleaved using 1 mg of PreScission to 500 mg of 3CLpro WT at 4 °C, before being flowed through a second Ni^2+^ column, and buffer-exchanged into crystallization buffer 1 (20 mM Tris-HCL pH 8.0, 150 mM NaCl, 1 mM TCEP).

#### 3CLpro WT crystallization

Platelike crystal clusters were produced by adding 10 µL of 3CLpro WT at 5 mg/mL to 10 µL of crystallization matrix (30% PEG 3350, 0.1 M Bis-tris propane pH 7.0) well solution in a hanging-drop vapor diffusion setup at 18 °C. Microseeds were generated from clusters using Seed-Beads (Hampton Research, USA). Single crystals were produced by setting up hanging drop crystallization wells at 18 °C with 10 µL of 3CLpro WT and 10 µL of 1:500 dilution microseeds, using 20% PEG 3350, 0.1 M Bis-tris propane pH 7.0 and 4 mg/mL 3CLpro WT protein concentration, and 20% PEG 3350/0.1 M Bis-tris pH 6.5, 5 mg/mL 3CLpro WT protein concentration.

#### 3CLpro C145S expression and purification

BL21(DE3) cells were transformed with pMCSG53 pDNA containing a 3CLpro C145S insert (provided by Andrzej Joachimiak) with a TEV-sensitive N-terminal His_6_ tag. Transformation, expression and harvesting was performed as with WT 3CLpro. Initial Ni^2+^ column purification was performed as with WT 3CLpro. Elution fractions with highest 3CLpro C145S concentration were combined and dialysed overnight against 2L of dialysis buffer (50 mM HEPES pH 7.2, 25 mM NaCl, 5% glycerol, 10 mM 2-mercaptoethanol). 48-hour TEV digestion was initiated by adding 25 ug of TEV per ug of 3CLpro C145S. The solution after cleavage reaction was passed through a second, 6 ml, pre-equilibrated nickel column. The flowthrough from this column was collected and buffer-exchanged using 10,000 MWCO centrifugal concentrators (EMD Millipore, USA) into crystallization buffer 2 (20 mM HEPES pH 7.2, 150 mM NaCl, 2 mM DTT).

#### 3CLpro C145S-NEMO binding

Human NEMO capped peptide (residues 226-235, acetyl-KLAQLQVAYH-amide, synthesized by Thermo Scientific) was dissolved to 2 mM in peptide buffer (20 mM HEPES, 150 mM NaCl, pH 7.2) and DMSO (5.67%). Peptides were then added to 3CLpro C145S at a 20-fold molar excess, before overnight incubation at 4 °C. The mixture was centrifuged to remove precipitants, and the supernatant was concentrated to 6.5 mg/mL for crystallization screens.

#### 3CLpro C145S crystallization

3CLpro C145S was incubated overnight with a 0.01-fold molar ratio of Human NEMO capped peptide and incubated overnight at 4 °C. No precipitation was observed. TOP96 (Rigaku Reagents, Japan), BCS (Molecular Dimensions, UK) and GRAS2 (Hampton Research, USA) screens were run using a Gryphon (Art Robbins Instruments, USA). 0.2 µL of protein solution at 6.5 mg/mL was added to 0.2 µL of crystallization matrix in a sitting-drop vapor diffusion setup at 18 °C. BCS screen condition A9 (0.1 M MES pH 6.5, 20 % v/v PEG Smear High) produced needle-like crystal clusters of 3CLpro C145S.

#### 3CLpro C145S-NEMO co-crystallization

TOP96 (Rigaku Reagents, Japan), BCS (Molecular Dimensions, UK) and GRAS2 (Hampton Research, USA) screens were run using a Gryphon (Art Robbins Instruments, USA). 0.2 µL of protein solution at 6.5 mg/mL was added to 0.2 µL of crystallization matrix in a sitting-drop vapor diffusion setup at 18 C. GRAS2 screen condition F11 (0.1 M Sodium phosphate dibasic dihydrate, pH 9.3, 10 mM Calcium chloride, 20% w/v Polyethylene glycol 3,350), yielded platelike single crystals. These crystals were isolated, cryoprotected by supplementation with 20% glycerol, mounted into ALS style, 0.05-0.1 mm loops (Hampton Research, USA) and flash frozen in liquid nitrogen.

#### Data collection and structural determination and refinement

Preliminary, cryogenic data collection was carried out at SSRL, using an Eiger X 16M detector, to confirm the presence of protein crystals. Cryogenic, X-ray crystallographic diffraction data of 3CLpro WT, C145S and NEMO-bound C145S was collected on beamline 12-2 at SSRL, using an Eiger X 16M detector. Data were processed using XDS. 3CLpro WT crystallized in the C2 space group, with one molecule in the asymmetric unit. Both 3CLpro C145S and the NEMO-bound 3CLpro C145S crystallized in the P1 space group with four protomers per asymmetric unit. The 3CLpro WT structure was phased in MOLREP ^50^ using the coordinates of 3CLpro bound to Telaprevir (PDB ID: 7K6D). For 3CLpro C145S and NEMO-bound 3CLpro C145S, molecular replacement was performed in MOLREP ^50^ using the coordinates of the 3CLpro WT structure solved in this study. Iterative refinement was performed manually in Coot ^51^ and REFMAC ^52^. The final rounds of refinement were performed by Phenix ^53^. Details of data collection and refinement parameters are provided in Supplementary Table S3.

### 5.3. Molecular dynamics simulations and binding affinity predictions

#### Preparation NEMO-bound models of 3CLpro from different betacoronaviruses

Chains A and B from the crystallographic structure described here were used as the starting configuration for the simulations of the NEMO-bound dimeric SARS-CoV-2 3CLpro. Along with that, structures of 3CLpro from SARS-CoV (PDB id 5B6O), MERS-CoV (PDB id 5C3N), and HKU1-CoV (PDB id 3D23) were used ^33, 54, 55^. After structural alignment, the coordinates from NEMO (aa. 227-234) from the X-ray structure of the NEMO-bound SARS-CoV-2 3CLpro C145S variant were used to build the models of NEMO-bound SARS-CoV, MERS-CoV, and HKU1-CoV 3CLpro dimers. In SARS-CoV-2 3CLpro, Val232 was replaced by Ala232 to also study the interactions in the binding site of 3CLpro with mouse NEMO_227-234_.

The models of complexes of 3CLpro with a long construct of human and mouse NEMO (aa. 190-270) were built prior to solving the crystallographic structure described in this study, so the coordinates of the 7-residue fragment of NEMO (residues 227-233) from the crystallographic structure of the porcine epidemic diarrhea virus (PEDV) 3CLpro bound to NEMO_227-233_ (PDB id 5ZQG) were used. We note that the NEMO-peptide in PEDV 3CLpro has a very similar conformation and position to what was found in SARS-CoV-2 3CLpro, displaying a NEMO_227- 233_ C_ɑ_/C_β_ RMSD of 0.9 Å between the two structures. NEMO N- and C-termini were capped with acetyl and N-methylamide, respectively. More details about system preparation are described in the Supplementary Methods.

#### Molecular dynamics simulations and conformation selection for binding affinity predictions

Different protocols of simulation were conducted for 3CLpro-NEMO_227-234_ and 3CLpro- NEMO_190-270_ systems, as the former was used both for traditional MD analysis and to select conformations for binding affinity predictions (Supplementary Methods). For the 3CLpro- NEMO_227-234_ systems, which were mostly generated *in silico*, we applied a strategy of MD simulations that aims structural refinement and sampling conformations that visit key interactions. Snapshots selected from these simulations were used as input for machine learning-based binding affinity prediction (MD/ML approach). Similarly to the method of protein structure refinement described in Heo et al.^56^, sampling was accelerated in a controlled manner by applying a fairly high temperature and weak position restraint potentials that compensate for the high thermal energy, which could drastically propagate the effects of any local molecular distortions or bad contacts. The applied position restraints are much weaker than the energy range of typical non-covalent interactions and conformational changes so that sampling is minimally biased while unrealistic states are easily escaped.

For the MD/ML approach, five independent conformational sampling runs were performed for 224 ns, recording coordinates every 40 ps. Supplementary Methods and Table S4 summarizes this MD-based protocol of sampling for conformation selection. The conformations generated from the restrained MD simulations were grouped via a root-mean-square deviation-based algorithm^57^. Considering only C_ɑ_ atoms of NEMO and residues in the Domains I and II of the *holo* chain of 3CLpro (aa. 16-198), the RMSD of atom-pair distances was computed applying the 1.0 Å cutoff as parameter for clustering. The interaction energy between 3CLpro and the NEMO- peptide was computed using the energy plugin within GROMACS. For each system, three structures with the lowest interaction energy were selected from different conformation clusters within the 20 most populated and used as input for ML-based binding affinity prediction.

In parallel, for classical molecular dynamics analysis, the equilibration phase was fully performed at 310.15 K and five independent production unrestrained runs of 112 ns were carried out. In the simulations of 3CLpro-NEMO_190-270_ systems, five production runs of 116 ns were performed with no position restraints.

#### Binding affinity from quantum mechanics calculations

The inclusion of quantum mechanics (QM) or machine learning (ML) in high-throughput drug screening to narrow down the list of promising inhibitor candidates has emerged as a very promising protocol. Recently, we illustrated an encouraging application of QM to refine the binding affinity of classical docking results of COVID-19 spike protein inhibitor drugs ^58^. In this work, we employed a similar strategy to evaluate the binding affinity of NEMO with different 3CLPro targets. For the QM-based prediction, we employed the linear-scaling fragment molecular orbital density-functional tight-binding (FMO-DFTB) ^59, 60^ method to predict binding affinities of NEMO with the four 3CLPro targets. In addition, the polarizable continuum model (PCM) was used to include solvent effects ^61^. The polarizable continuum model (PCM) was used to describe interactions with solvent, and the empirical D3 dispersion correction was used to improve the accuracy in describing non-covalent interactions. For the PCM calculations, the cavity was calculated using newly calibrated atomic radii to improve accuracy in predicting solvation free energy of DFTB in an aqueous solvent. The details of the atomic radii optimization will be published elsewhere. For each target, the FMO-DFTB/PCM method was used to re-optimize structures of NEMO model systems in bound and unbound states, while the target’s structure was fixed. Partial re-optimization of the target did not significantly alter the result. The internal binding energy is defined as the difference in internal energy between the complex and its corresponding unbound NEMO and unbound 3CLPro.

#### Binding affinity from machine learning-based rescoring

Machine-learned models dedicated to predicting binding affinities (Demerdash, 2021, *in press*) were trained on a large database of protein-small molecule complexes with both experimental binding affinities and X-ray crystal complexes known as PDBBIND ^62^, followed by testing on an independent data set, the CASF-2013 benchmark ^62^, that was not used in the training. For each complex a set of seven features were calculated, where each feature is itself a predicted affinity or free energy calculated under very different physical assumptions (Supplementary Methods).

Models were trained using either support-vector machines, gradient-boosted trees, or random forest. For each of these 3 methods, models were trained using normalized, or scaled, features. For the gradient-boosted tree and random forest method, models were additionally trained on the raw, unscaled features. For each *betacoronavirus* 3CLpro, the models were applied to the energy minimized complexes and to the set of MD-generated conformers. Separately, both the full set of MD conformers as well as the three conformers with the lowest molecular mechanics (MM) energy of interaction were considered for each system. The average predicted affinity was obtained, considering either the full set of MD conformers or just the three with the lowest MM energy of interaction (Figure 4b). Noticeably, the ML models applied to energy minimized complexes (conformers prior to MD simulations) did not provide consistent results among the rankings. These structures have appreciably higher MM energy than the MD structures, suggesting that the consistent rankings of the MD structures reflect more native-like structures and that the MD protocol of confirmation selection is effective to account for the conformational flexibility of the receptor.

#### Molecular re-docking of 3CLpro C-terminus

Chains A and B from the crystallographic structure of SARS-CoV-2 3CLpro-NEMO described in this study were used to estimate the contributions of local interactions to the binding affinity of the C-terminus to the dimer interfacial site. The C-terminus of chain B seats between Domains I in the dimer formed with chain A. Chains A and D were used as receptor and ligand, respectively, to estimate the contributions of local interactions to the binding affinity of the C-terminus to a catalytic site in a neighboring dimer. For the calculations of C-terminus at the interfacial and catalytic binding sites, a peptide comprising residues 302 to 306 was defined from the C-terminus of chain B and D, respectively, detaching it from the rest of the protein by deleting residues 300 and 301. N- and C-termini of these peptides were capped with acetyl and

N-methylamide, respectively. Molecular re-docking of the C-terminus peptides was performed using AutoDock Vina ^63^. All bonds of the peptide were kept rigid as the goal was to preserve the initial conformation and compute the binding free energy using the AutoDock Vina scoring function.

