## Supplementary Methods, Tables S1-S4, and Figures S1-S6 for "Structural and functional characterization of NEMO cleavage by SARS-CoV-2 3CLpro"

Content:

- Supplementary Methods
- Supplementary Tables S1-S4
- Supplementary Figures S1-S6

**Supplementary Methods**

### Molecular dynamics simulations and binding affinity predictions

*Preparation of systems for simulations*

Structural alignments were done using the MultiSeq[1] plugin within VMD[2]. Missing residues were added using coordinates of backbone atoms from the structure of SARS-CoV-2 3CLpro C145S variant. CHARMM-GUI[3] was used to model the missing side chains, to replace mutated residues by native residues, and to prepare simulation systems and inputs. [1] Structural water molecules were kept in the built systems.

Schrödinger Bioluminate Version 2019-4 [4] was used to construct the model of complexes of 3CLpro with a long construct of human NEMO (aa. 190-270). Using the Bioluminate's build function and the coiled coil structure of NEMO from PDB id 6MI3 as template [5], the NEMO fragment in the active site of 3CLpro was extended to generate a construct comprising residues 190-270 (human $NEMO_{190-270}$). Val232 was replaced by Ala232 to study the interactions in the binding site of 3CLpro with mouse $NEMO_{190-270}$.

The simulation boxes were generated by defining a solvation layer of 10 Å minimum thickness around the protein complex. 0.15 M KCL was used to establish electroneutrality. Protonation states for all systems were defined based on the neutron crystallographic structure of SARS-CoV-2 3CLpro (PDB id 7JUN) [6], which informs fine atomic details at a near-physiological pH (6.6).

*Molecular dynamics simulations for classical analysis and for conformation selection*

GROMACS-2020 [7] was used to run all simulations on the Summit supercomputer at the Oak Ridge Leadership Computing Facility. For all systems, initial energy minimization was performed with steepest descent for 5000 steps. The particle mesh Ewald method[8] was used for the treatment of periodic electrostatic interactions, using a cutoff distance of 12 Å. The Lennard–Jones potential was smoothed over the cutoff range of 10–12 Å. LINCS[9] was used to constrain all bonds involving hydrogen atoms. CHARMM36m[10–12] and TIP3P[13] force fields were used to describe protein and water molecules, respectively. Hydrogen mass repartitioning[14] was applied to the full 3CLpro-NEMO$_{227\text{-}234}$ systems to accelerate sampling as it allows using a 4 fs integration time step.

In the simulations for the MD/ML approach applied to the 3CLpro-NEMO$_{227\text{-}234}$ systems, velocity Langevin dynamics was performed using a friction constant of 1 ps$^{-1}$. An initial equilibration phase of about 5 ns for SARS-CoV-2 3CLpro and 24 ns for the other *betacoronaviruses* 3CLpro was conducted for a gradual increase in temperature up to 320.15 K and gradual release of position restraint harmonic potentials applied to $C_\alpha$ atoms of 3CLpro and the NEMO-peptide. To allow the box size to adjust, most of the equilibration phase was conducted in the NpT ensemble, using the Berendsen barostat[15] applying a compressibility of 4.5 X 10$^{-5}$ bar$^{-1}$ and a time constant of 1.0 ps. In the last 1 ns of equilibration, the barostat was switched off and the next steps were carried out in the NVT ensemble. In the final step of equilibration, the atomic velocities were reinitialized independently from a Maxwell-Boltzmann distribution using random numbers of seed.

Following equilibration, five independent restrained MD simulations were performed for each system for conformational sampling and selection. Flat-bottom harmonic restraint potentials were applied to all $C_\alpha$ atoms of 3CLpro, using a force constant of 0.25 kcal/mol/Å$^2$ and a flat-bottom width of 4 Å. Harmonic position restrain potential were kept at the $C_\alpha$ atoms of Gln229, Leu230, and Gln231 in NEMO using a force constant of 0.05 kcal/mol/Å$^2$.

In the simulations of 3CLpro-NEMO$_{190\text{-}270}$ systems, a long equilibration phase at 310.15 K was performed for about 120 ns with a gradual release of position restraints applied to the backbone atoms of 3CLpro and NEMO via harmonic potentials. During the last 76 ns, weak position restraints were applied only to the backbone atoms of residues of NEMO at the binding site of 3CLpro (aa. 227-233) using a force constant of 0.0125 kcal/mol/Å$^2$. Temperature control at 310.15 K was performed with the Nosé–Hoover thermostat using a time constant of 1.0 ps[16].The Parrinello-Rahman barostat with coupling constant of 1.0 ps was used with 1 atm as reference[17]. In the final step of equilibration, the atomic velocities were reinitialized independently and five production runs of 116 ns were performed with no position restraints.

Visual Molecular Dynamics software suite (VMD, version 1.9.4a48)[18] was used for visual analysis. The Hbonds plugin in VMD was used to compute statistics of hydrogen bond interactions. The geometric criteria adopted are a cutoff of 3.0 Å for donor-acceptor distance and 20° for acceptor-donor-H angle. VMD was also used to compute distances and $C_\alpha$ RMSD of selected atoms to a reference structure. Average number of contacts was computed using the mindist utility of GROMACS and a distance cutoff between $C_\alpha$ atoms of 8 Å. The main contacts were identified using the Timeline plugin in VMD. Grace was used for the timeline plots (https://plasma-gate.weizmann.ac.il/Grace/).

*Binding affinity predictions*

All FMO-DFTB/PCM calculations were carried out using the GAMESS program.[19] In the machine learning-based rescoring, the affinities/free energies corresponding to the seven features were obtained with the following freely available software: X-Score [20], DSX [21], RF-Score-VS [22], Deltavina [23], RF-Score-3 [24] , GalaxyDockBP2 [25], and Cyscore [26]. Autodocktools [27] was used to prepare the inputs for molecular re-docking.

**Supplementary Tables**

**Supplementary Table S1. Hydrogen bond interactions computed from molecular dynamics simulations of the SARS-CoV-2 3CLpro-NEMO bound complex**. The average of occurrence (AVG) and its standard deviation (SD) are shown as percentage of simulation time steps in five MD runs of 100 ns each. Corresponding pairs of residues as well as atom name are identified ([residue]_[atom]_[side or main chain]).

| **3CLpro** | **NEMO** | **AVG** | **SD** |
|---|---|---|---|
| Glu166_N/O_main | Gln229_O/N_main | 93 | 2 |
| Thr26_N/O_main | Ala233_O/N_main | 88 | 4 |
| Thr190_O_main | Ala228_N_main | 63 | 15 |
| Gly143_N_main | Gln231_O_main | 41 | 8 |
| His164_O_main | Gln231_N_main | 36 | 20 |
| Gln189_N_side | Ala228_O_main | 30 | 8 |
| His163_Nε2_side | Gln231_O_side | 33 | 16 |
| Cys145_N_main | Gln231_O_main | 30 | 3 |
| Asn142_N_side | Val232_O_main | 21 | 4 |

**Supplementary Table S2. Comparisons of 3CLpro recognition site at Gln231 in NEMO (aa. 226-235) from multiple animal species.** Light green highlights the reference sequence, hNEMO, and light purple, species exhibiting a different motif.

| Order | Organism | Cleavage site $NEMO_{226-235}$ | Reference NCBI |
|---|---|---|---|
| Primates | **Human** *Homo sapiens* | KLAQLQVAYH | NP_001093327.1 |
| | **Chimpanzee** *Pan troglodytes* | KLAQLQVAYH | XP_016800099.2 |
| | **Rhesus monkey** *Macaca mulatta* | KLAQLQVAYH | XP_001095498.2 |
| | **Marmoset** *Callithrix jacchus* | KLAQLQVAYH | XP_008988365.2 |
| | **Capuchin** *Sapajus apella* | KLAQLQVAYH | XP_032129057.1 |
| | **Crab-eating macaque** *Macaca fascicularis* | KLAQLQVAYH | XP_005595059.1 |
| Rodents | **Norway rat** *Rattus norvegicus* | KLAQLQAAYH | NP_954534.1 |
| | **Thirteen-lined ground squirrel** *Ictidomys tridecemlineatus* | KLAQLQVAYH | XP_040143177.1 |
| | **Chinese hamster** *Cricetulus griseus* | KLAQLQAAYH | XP_016836062.1 |
| | **Mouse** *Mus musculus* | KLAQLQAAYH | NP_001154895.1 |
| Carnivores | **Domestic ferret** *Mustela putorius furo* | KLAQLQVAYH | XP_012905560.1 |
| | **Stoat** *Mustela erminea* | KLAQLQVAYH | XP_032187502.1 |
| | **Domestic cat** *Felis catus* | KLAQLQVAYH | XP_004001080.1 |
| | **Tiger** *Panthera tigris* | KLAQLQVAYH | XP_042830262.1 |
| | **Dog** *Canis lupus familiaris* | KLAQLQVAYH | XP_038307227.1 |
| Even-toed ungulates | **Horse** *Equus caballus* | KLAQLQVAYH | NP_001271462.1 |
| | **Wild boar** *Sus scrofa* | KLAQLQVAYH | NP_001106524.1 |
| | **Dromedary** *Camelus dromedarius* | KLAQLQVAYH | XP_031302174.1 |
| Rabbit | **Rabbit** *Oryctolagus cuniculus* | KLAQLQVAYH | NP_001164965.1 |

**Supplementary Table S2 (cont.). Comparisons of 3CLpro recognition site at Gln231 in NEMO (aa. 226-235) from multiple animal species.** Light green highlights the reference sequence, hNEMO, and light purple, species exhibiting a different motif.

| Order | Organism | Cleavage site<br>$NEMO_{226-235}$ | Reference<br>NCBI |
|---|---|---|---|
| Bats | **Common vampire**<br>*Desmodus rotundus* | KLAQLQVAYH | XP_024406864.1 |
| | **David's myotis**<br>*Myotis davidii* | KLAQLQAAYH | XP_006754835.1 |
| | **Great roundleaf**<br>*Hipposideros armiger* | KLAQLQVAYH | XP_019489218.1 |
| | **Black flying fox**<br>*Pteropus alecto* | KLAQLQVAYH | XP_006904856.1 |
| | **Natal long-fingered**<br>*Miniopterus natalensis* | KLAQLQVAYH | XP_016077428.1 |
| | **Brandt's**<br>*Myotis brandtii* | KLAQLQAAYH | XP_005886157.1 |
| | **Large flying fox**<br>*Pteropus vampyrus* | KLAQLQVAYH | XP_039722945.1 |
| | **Egyptian rousette**<br>*Rousettus aegyptiacus* | KLAQLQVAYH | XP_015979392.2 |
| | **Greater horseshoe**<br>*Rhinolophus ferrumequinum* | KLAQLQVAYH | XP_032959395.1 |
| | **Pale spear-nosed**<br>*Phyllostomus discolor* | KLAQLQVAYH | XP_028378091.1 |
| | **Big brown**<br>*Eptesicus fuscus* | KLAQLQAAYH | XP_027991937.1 |
| Pholidota | **Pangolin**<br>*Manis javanica* | KLAQLQVAYH | XP_017521897.1 |

**Supplementary Table S3. Crystallographic parameters, data collection and refinement statistics.**

| | 3CLpro-NEMO | 3CLpro C145S | 3CLpro WT |
|---|---|---|---|
| **Crystallographic parameters** | | | |
| Space group | P1 | P1 | C2 |
| Unit-cell dimensions | 63.39Å, 67.71Å, 77.84Å<br>102.5˚, 89.9˚,107.4˚ | 61.59Å, 67.51Å, 77.65Å<br>102.1˚, 89.3˚,106.4˚ | 114.17Å, 53.44Å, 44.87Å<br>90.0˚, 103.0˚, 90.0˚ |
| **Data collection statistics** | | | |
| Resolution limits (Å) | 38.1 – 2.14 | 38.05-2.45 | 38.9-1.45 |
| No: of observed reflections | 335783 | 339666 | 535143 |
| No: of unique reflections | 64759 | 41750 | 46003 |
| Completeness overall/outer shell | 97.5/96.3 | 96.9/93.6 | 98.4/98.4 |
| CC1/2 (overall/outer shell) | 99.3/61.5 | 99.4/58.6 | 99.9/68.1 |
| $R_{sym}^{a}$ (%)<br>overall/outer shell & os I/σ | 18.4/95.5 & 1.8 | 28.3/108.0 & 1.6 | 7.5/156.6 & 1.8 |
| **Refinement statistics** | | | |
| Resolution limits (Å) | 38.1-2.14 | 38.05-2.45 | 38.9-1.45 |
| Number of reflections/% | 61520/97.6 | 39661/96.9 | 43702/98.4 |
| (\|F\|>2σ\|F\|) | 3238 | 2088 | 2301 |
| Reflections used for $R_{free}$ | | | |
| $R_{factor}^{b}$ (%) | 21.4 | 19.4 | 15.5 |
| $R_{free}$ (%) | 30.9 | 27.3 | 20.0 |
| Model contents/average B(Å$^2$) | | | |
| Protein atoms | 9439/42.9 | 9421/51.6 | 2403/33.4 |
| Peptide | 154/46.8 | 0 | 0 |
| Water molecules | 193/37.6 | 39/34.0 | 227/39.6 |
| RMS deviations | | | |
| Bond length (Å) | 0.010 | 0.008 | 0.008 |
| Bond angle (°) | 1.50 | 1.64 | 1.47 |
| Ramachandran (analyzed/outliers) | 1225/6 | 1222/18 | 305/0 |

**Supplementary Table S4. Molecular dynamics-based protocol of sampling for conformation selection of 3CLpro-NEMO$_{227\text{-}234}$.** In the equilibration phases, there is a gradual change in temperature, in time step, and in the position restraint potentials. Atom velocities are re-distributed using five different seed numbers to initiate equilibration_6. With that, five independent trajectories are generated for conformational sampling. Applied potentials are either simple harmonic restraints (H) or flat-bottom potentials (FB). In NEMO$_{227\text{-}234}$, position restraints were applied to $C_\alpha$ atoms of residues 229-231.

| Simulation phases | sition Restraints | | | | Time Step | Total Time | Temperature |
|---|---|---|---|---|---|---|---|
| | 3CLpro - $C_\alpha$ | Type | 3CLpro - $C_\alpha$ | Type | | | |
| **Equilibration_1** | 0.50 | H | 0.25 | H | 1 fs | 125 ps | 100.15 K |
| **Equilibration_2** | 0.25 | H | 0.25 | H | 2 fs | 500 ps | 200.15 K |
| **Equilibration_3** | 0.14 | H | 0.25 | H | 2 fs | 500 ps | 250.15 K |
| **Equilibration_4** | 0.05 | H | 0.25 | H | 2 fs | 2 ns | 300.15 K |
| **Equilibration_5** | 0.05 | H | 0.14 | H | 4 fs | 8 ns | 310.15 K |
| **Equilibration_6** | 0.14 | FB | 0.10 | H | 4 fs | 1 ns | 320.15 K |
| **Sampling** | 0.25 | FB | 0.10 | H | 4 fs | 120 ns (x 5) | 320.15 K |

### Supplementary Figures

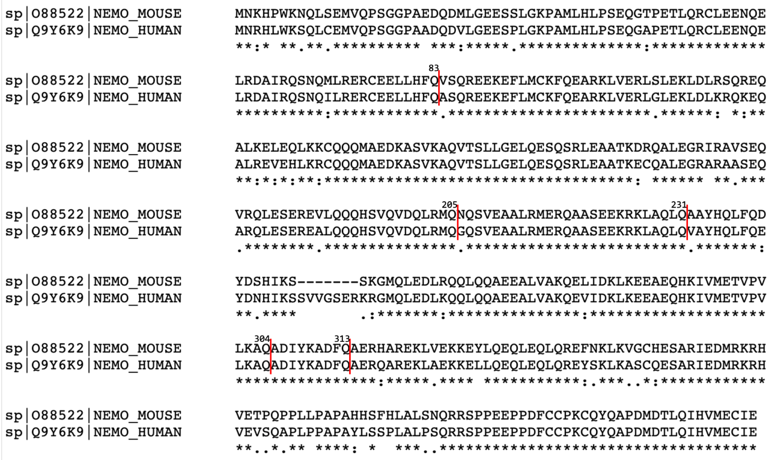


**Supplementary Figure S1. Predicted 3CLpro cleavage sites in mouse and human NEMO.** Aligned sequences of mouse (*Mus musculus*) and human NEMO are shown. The five predicted 3CLpro recognition sites at Gln83, Gln205, Gln231, Gln304, and Gln313 are indicated with the red lines.

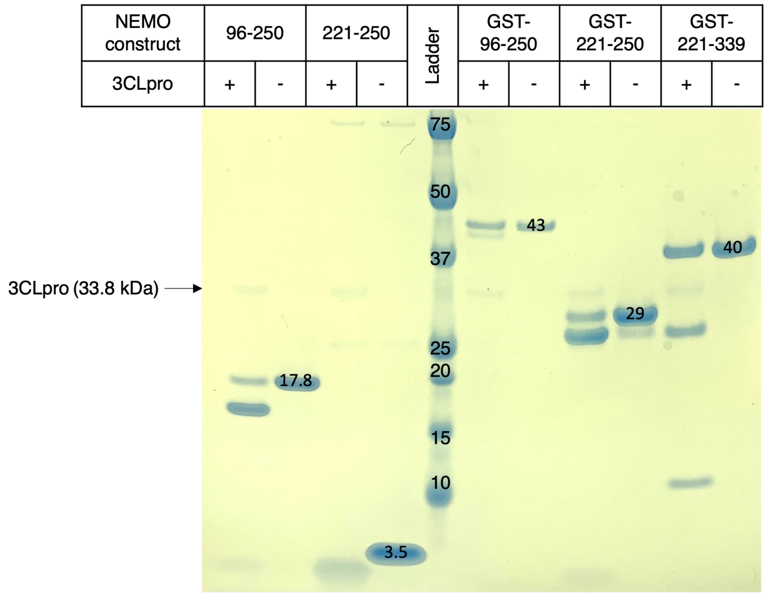


**Supplementary Figure S2. 3CLpro cleaves mouse NEMO.** SDS-PAGE following incubation of five truncations of mouse NEMO (0.5 mg/mL) with and without 3CLpro (250 nM). Proteolysis products are consistent with a single cleavage site following Gln231.

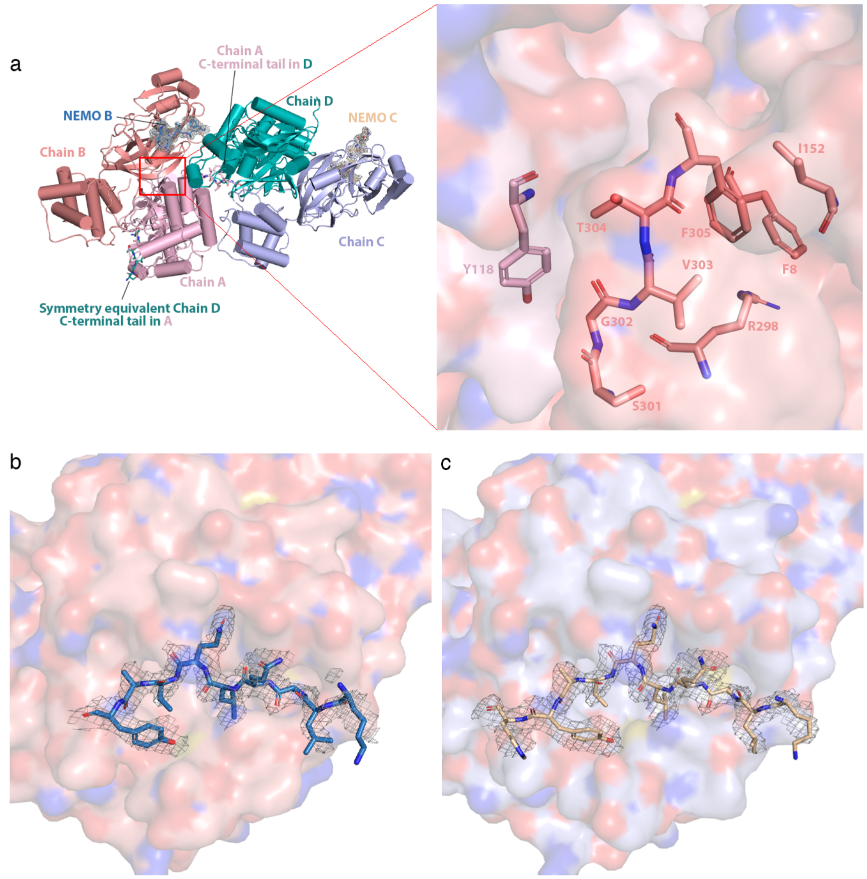


**Supplementary Figure S3. Structure of NEMO-bound C145S variant asymmetric unit, the interfacial C-terminal-binding site and NEMO peptide densities**. **a)** Chains A, B, C, and D are colored light pink, salmon, light blue, and teal, respectively. The C-terminal tail of chain A is depicted as sticks and binds into the substrate-binding site of chain D. NEMO bound into chain B (NEMO B) is displayed as sticks and colored dark blue. NEMO C is displayed as sticks and colored wheat. A mesh of density is displayed around NEMO B and NEMO C, in light grey. Inset, the C-terminal tail of chain B is depicted as sticks in salmon. Residues Ser301 to Phe306 are labelled. The surfaces of chains A and B are shown in pink and salmon, respectively, and both colored by atom type, where oxygens are red, nitrogens are dark blue, carbons are pink or salmon and sulfurs are yellow. Residues in chain A that interact with the C-terminal tail of chain B are portrayed as sticks and labelled. **b)** Electron density around NEMO B. **c)** Electron density around NEMO C.

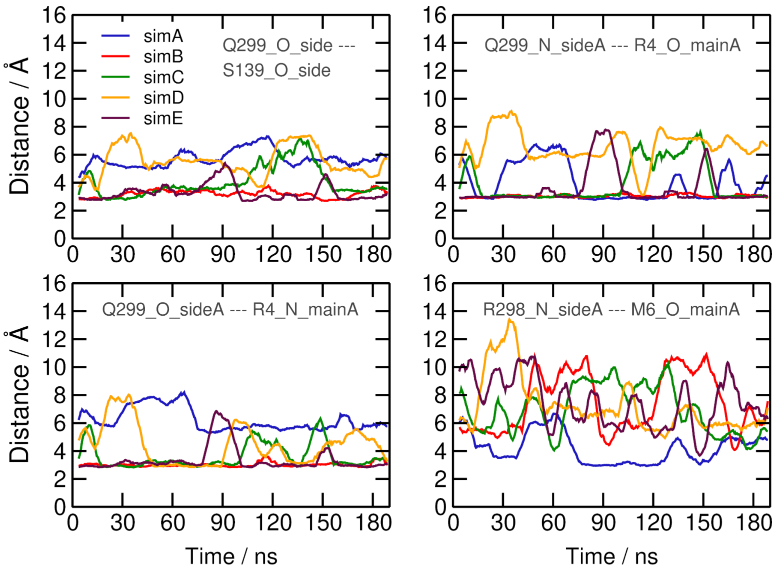


**Supplementary Figure S4. Time evolution of key distances involving the C-terminal helix of SARS-CoV-2 3CLpro in molecular dynamics simulations.** Distances involving Arg298 and Gln299 in chain A of the structure of NEMO-bound 3CLpro computed from five MD trajectories (simA-E). Corresponding pairs of residues as well as atom name and protein chain are identified in each panel ([residue]_[atom]_[side or main chain][chain]).

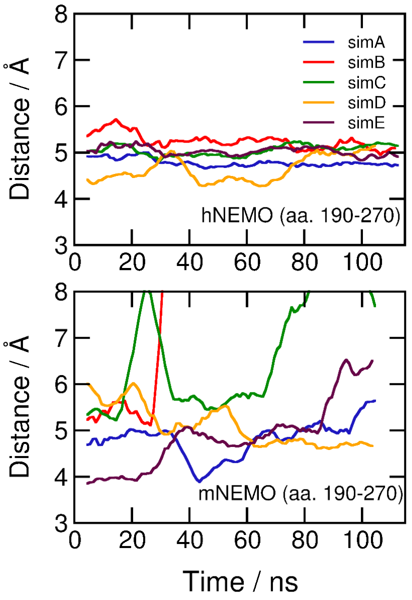


**Supplementary Figure S5. Time evolution of the distance between the catalytic S⁻ in 3CLpro Cys145 and the carbonyl C of Gln231 in human and mouse NEMO$_{190\text{-}270}$ computed from MD simulations.** Distances computed from five independent molecular dynamics simulations (simA-E).

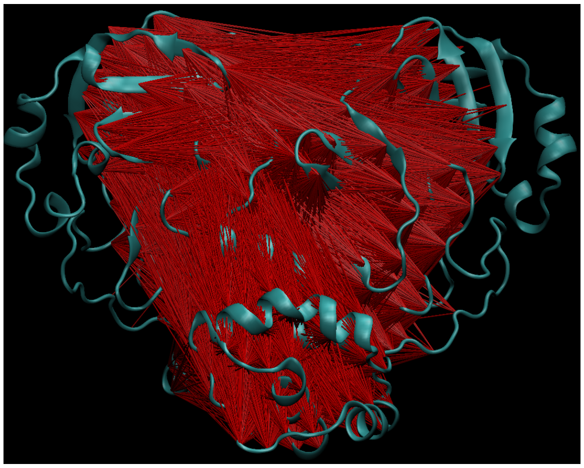

**Supplementary Figure S6. Essential $C_{\alpha}$ cross-correlation analysis of SARS-CoV-2 3CLpro**. Representative structural network of the 35707 pairs of residues that exhibit negative correlations (red sticks) with modulus greater than 0.85 identified in the MD trajectories of the *apo* SARS-CoV-2 3CLpro dimer (cyan).
